# Microstructural Analysis of Language Pathways’ Development in Term Neonates

**DOI:** 10.64898/2026.11.23.767641

**Authors:** Ali Kazemi, Roxana Namiranian, Hamid Abrishami Moghaddam

## Abstract

Understanding the early development of language pathways is essential for elucidating how neonatal brain organization supports the acquisition of linguistic skills. In this study, we investigated microstructural changes in term human neonates scanned at postmenstrual ages (PMA) of 37 to 44 weeks. Using diffusion MRI data from 310 healthy infants in the developing Human Connectome Project, we employed q-space diffeomorphic reconstruction to map major language tracts involved in language processing. Fractional anisotropy, mean diffusivity, and other diffusion-based biomarkers were quantified in pathways such as the arcuate fasciculus, extreme capsule, and inferior fronto-occipital fasciculus. Our results revealed significant weekly differences in microstructural integrity across these tracts, with particularly robust age-related reductions in mean and radial diffusivity. In addition, we observed leftward asymmetries in multiple language pathways, suggesting that hemispheric specialization emerges by this early postnatal stage. Although PMA at scan predicted the most pronounced shifts in diffusion metrics, sex showed minimal impact on developmental trajectories. These findings highlight the rapid and tract-specific organization of language-relevant connections during the first weeks of life, providing normative data against which atypical patterns can be evaluated. More broadly, this work offers new insights into the foundation of language circuitry in neonates, informing future studies of neurodevelopmental disorders and potential early-life interventions.

## 1. Introduction

The cognitive process of language acquisition represents a primary focus of investigation in developmental neuroscience. A critical phase of this development occurs during the prenatal and postnatal periods— epochs that remain significantly understudied compared to childhood and adulthood. Furthermore, experimental auditory and language-related stimuli [1], [2] do not require active or deliberate behavioral responses from the infant subject. Because the maturation of underlying neural structures heavily dictates early language development, establishing normative developmental trajectories of language-associated white matter tracts in term neonates is essential. Such normative data will enable neuroscientists to better understand the functional development of language-related regions and design more robust developmental experiments.

Language pathway development is a prolonged process which commences at the seventeenth post-conceptional week (PCW) [3], [4], undergoes rapid alteration in the first two years of life, and continues until early adulthood [5]–[8]. Any deviation from normal developmental patterns may result in disorders such as aphasia, stuttering, alexia, dyslexia, amusia, and apraxia [9]. With normative pre- and postnatal developmental patterns in hand, language-related brain anomalies could be diagnosed and treated at an early stage of development.

The perisylvian cortex is the primary neural region responsible for language processing. This region encompasses the cortical areas surrounding the lateral sulcus (Sylvian fissure), including Broca’s area (composed of the pars opercularis and pars triangularis of the inferior frontal gyrus) and Wernicke’s area (the posterior portion of the superior temporal gyrus) [10]. Over the last several decades, our understanding of the microstructural underpinnings of language has significantly evolved. While the arcuate fasciculus (AF) was historically considered the sole pathway dedicated to transmitting language-related signals, contemporary models of language computation emphasize a more spatially distributed network. Alongside the AF, modern dual-stream and multi-tract models incorporate several other critical pathways, including the superior longitudinal fasciculus (SLF), inferior fronto-occipital fasciculus (IFOF), uncinate fasciculus (UF), frontal aslant tract (FAT), extreme capsule (EC), inferior longitudinal fasciculus (ILF), vertical occipital fasciculus (VOF), and middle longitudinal fasciculus (MLF) [7], [11]–[14]. These language tracts establish complex networks that connect various cortical regions across the frontal, temporal, occipital, and parietal lobes. Functionally, these pathways support higher-order processes essential for phonology, semantics, literacy, visuo-spatial word recognition, and motor-vocal coordination tasks. Supplementary Table S1 outlines the anatomical and functional characteristics of these tracts in detail.

In recent years, noninvasive brain imaging technologies have been widely used for examining the anatomical structure of human brain pathways. Diffusion tensor imaging (DTI) can depict the maturation of pathways from the fetal period [4], [15], [16] to adulthood. Diffusion metrics suggest that during the growth of the human brain, fractional anisotropy (FA) increases, and mean diffusivity (MD) decreases [17]–[19]. Nevertheless, while DTI measures are sensitive markers of white matter maturity and integrity, they are nonspecific and assess many aspects of white matter microstructure [20], [21]. Furthermore, DTI measures are confounded by factors such as crossing or kissing fiber populations, partial volume effects, and processing pipeline biases [21], [22]. Advanced diffusion imaging methodologies like high angular resolution diffusion imaging (HARDI) [23] were developed to overcome the limitations of DTI and provide new anisotropy measures beyond the classical FA. Developmental changes in intra-axis space, inter-axis space, and cerebral CSF can be indicated using q-ball reconstruction methods such as GQI [24] and QSDR [25]. The well-known metrics based on these methodologies are quantitative anisotropy (QA), isotropic diffusion (ISO), and restricted diffusion index (RDI).

Language pathways development starts from the third trimester and continues until adulthood. Although by the 35th week of GA, all the language tracts are connected to their neuronal endings, myelination has not yet begun [3]. Nevertheless, functional examination of brain development illustrates that in 3-month-old infants, speech stimuli activate temporal regions [2]. Each language fiber develops at a different speed, and maturation of pathways such as AF and SLF continues into early adulthood [6]. Ventral linguistic pathways (i.e., MLF, IFOF, UF, EC, and ILF) develop faster than dorsal linguistic pathways (AF and SLF) [7]. The fast development of ventral pathways is reported in 6-22 weeks after birth using 21 healthy infants [7]. In infants born preterm and scanned up to five months, the IFOF and ILF already show an anterior–posterior developmental gradient [26]. Due to the diversity in maturation rates of the brain’s fibers, more studies using sufficient numbers of subjects are needed to cover the vital periods of development with appropriate time resolution to map the maturation pattern of all language-related fibers.

From early on, perisylvian regions are anatomically asymmetric [27]–[29], suggesting their functional lateralization in language processing prior to language exposure. Specifically, AF has strong laterality that continues until adulthood. Nevertheless, functional lateralization of language changes with age [30], and in the first postnatal months the lateralization of the major tracts varies with age and between tracts [31]; asymmetry of white-matter microstructure is already detectable in infants [32]. AF in each hemisphere is dedicated to a different task. The left AF is associated with syntactic processing [33], [34]; and the right AF plays a role in prosody and semantics [34]. Surprisingly, verbal short-term memory and language processing are mediated by the same areas of inferior frontal and posterior temporal sectors in the left hemisphere [35].

Evidence for sex differences in brain and language development is limited. Sex differences in brain structure or function do not necessarily lead to differences in language task performance [36]; in infants, sex differences in white-matter microstructure have not been detected [32]. Delineating sex hormones in neonates and infants [37] demonstrates that the postnatal sex hormone surge might be viewed as one factor that influences later language development, independent of gender.

Scrutinizing anthropometric indicators in commissural and projection fibers in full-term neonates [38], [39] has indicated that subjects with higher birth indicators such as GA, birth weight, and crown-heel length may show greater brain microstructural maturation. Among the four birth indicators examined, GA showed the strongest association with WM maturation.

The majority of brain development research concentrates on global pathway maturation, with a particular emphasis on prominent structures such as the corpus callosum and AF. Among language tracts, AF is the only tract that is well-examined from early life to adulthood. Furthermore, the current body of research investigating brain pathway development remains limited. Given the rapid developmental changes in the neonatal brain and documented substantial growth, achieving adequate temporal resolution in this population is challenging. Furthermore, findings from these studies are derived from conventional diffusion tensor imaging (DTI) techniques, which, as previously discussed, encounter significant challenges, particularly in neonatal brain imaging. Consequently, a normative model for language pathway development in neonates remains elusive.

In this study, we present a weekly model for the development of language pathways. Normative atlases of diffusion metrics [40] can help radiologists identify early divergence from normal development. The image quality and acquisition of multi-shell HARDI data maximized the DTI tissue contrast [41], which allowed us to use advanced analysis techniques. We examine the alteration in anatomical structures of the brain via both DTI metrics, such as FA and MD, and measurements from QSDR reconstruction methodology, such as QA and RDI. Moreover, the dHCP dataset has provided anthropometrics such as birth weight and head circumference, which are used to better understand the development of the neonatal brain.

## 2. Material and Methods

The processing and analysis pipeline is summarized in Fig. 1.

**Fig 1.**
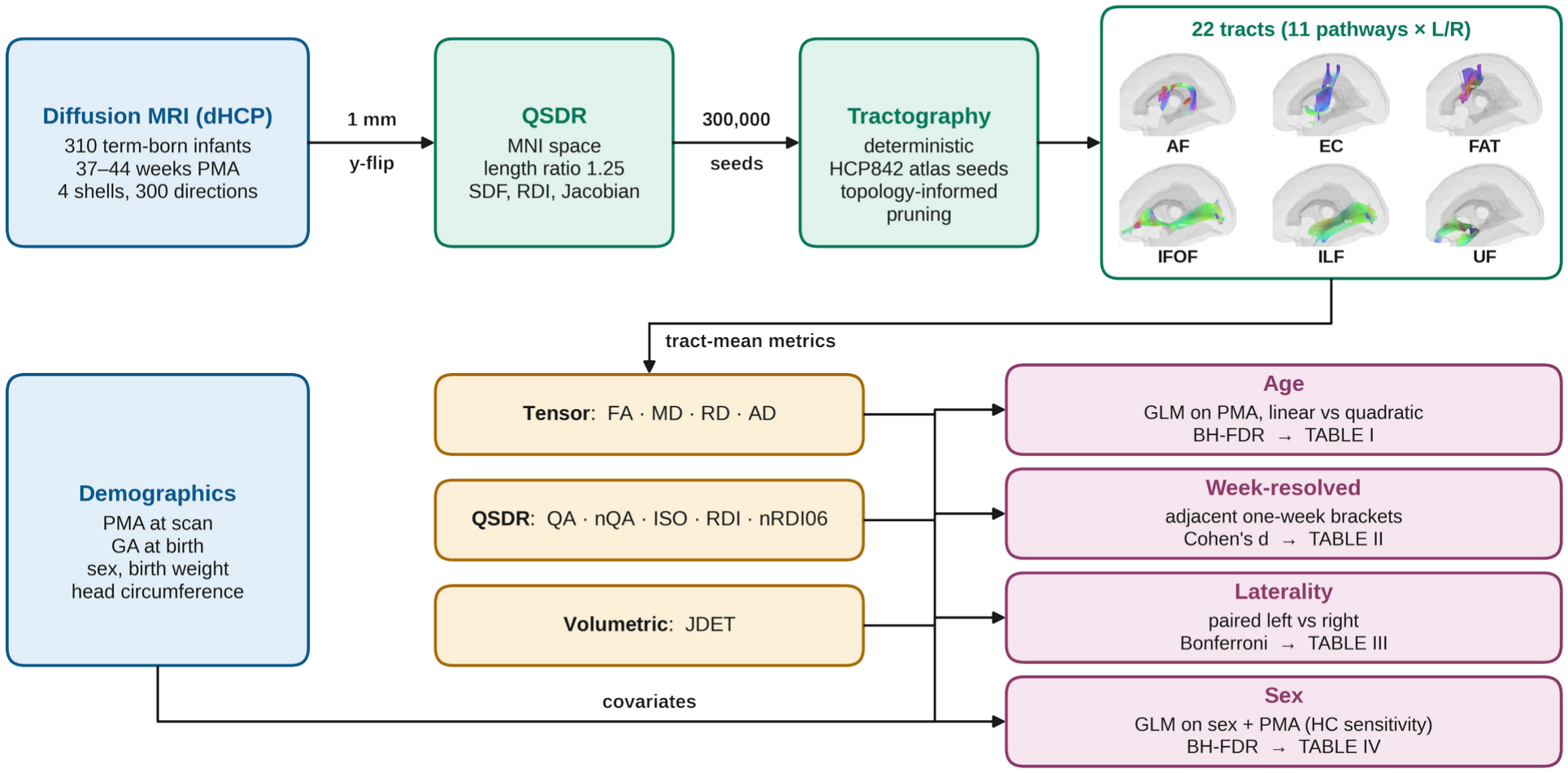
Overview of the processing and analysis pipeline. Diffusion MRI of 310 term-born dHCP infants was resampled to 1 mm and reconstructed in MNI space with QSDR; 22 language tracts (11 pathways, left and right; six shown on DSI Studio’s dHCP neonatal template) were reconstructed by atlas-seeded deterministic tractography with topology-informed pruning. Tract-mean tensor, QSDR and volumetric metrics, with the demographic covariates, entered four analyses, each reported in one table. HC, head circumference.

### 2.1 Subjects

We used structural and diffusion MRI (multishell HARDI) data from the second release of the dHCP (www.developingconnectome.org). Of 368 term-born neonates (GA at birth ≥ 37 weeks), we excluded infants with missing anthropometric information, such as age or head circumference, and infants with corrupted diffusion images. Where a scan session had been reconstructed more than once, we kept the reconstruction made with the study’s parameters (diffusion sampling length ratio 1.25) and the best template registration. For the one infant scanned twice, we kept the earlier scan, so that every infant contributed a single observation. The final sample comprised 310 infants (167 male, 143 female), whose age distribution is shown in Supplementary Fig. S1 and Supplementary Table S2. The infants were born at a median GA of 40.1 weeks (interquartile range 39.0–40.9) and scanned at a median postmenstrual age (PMA) of 40.9 weeks (39.6–41.9).

Images were acquired on a 3 T Philips Achieva scanner with a system dedicated to neonatal brain imaging [42]. Diffusion data were collected over four b-value shells (0, 400, 1000 and 2600 s/mm ^2^) with 300 encoding directions (20, 64, 88 and 128 per shell) [41]. The in-plane resolution was 1.17 mm, and the slice thickness was 3 mm with a 1.5 mm overlap. All images had been preprocessed with the automated dHCP neonatal diffusion pipeline, which includes off-resonance field estimation, distortion correction, super-resolution reconstruction and registration to standard space [41], [43].

### 2.2 Data Processing

The diffusion-weighted images were resampled to 1 mm isotropic resolution, and the y-axis was flipped to match the orientation of the Montreal Neurological Institute (MNI) space. The data were reconstructed in MNI space with QSDR [25] to obtain the spin distribution function [24], using a diffusion sampling length ratio of 1.25 (with one exception, described in Section 2.5). Restricted diffusion was quantified with restricted diffusion imaging [44], and the Jacobian determinant of the subject-to-template transformation was used as a measure of local volume [45], [46].

Fiber tracking used a deterministic algorithm [47]. Each tract was seeded from its white-matter region in the HCP842 tractography atlas [48], after nonlinear registration of the atlas to the subject. The anisotropy threshold was 0.04, the angular threshold 60°, and the step size 0.5 mm; streamlines shorter than 15 mm or longer than 150 mm were discarded. For each of the 22 tracts, 300,000 seeds were placed, and topology-informed pruning [49] was applied for 20 iterations to remove spurious connections. Tract terminations were checked visually to confirm that they reached the expected cortical regions, which are shown in Fig 2. For illustration, the 22 tracts are shown on the dHCP neonatal template of DSI Studio in Supplementary Fig. S2. After duplicate reconstructions were removed, ten metrics were extracted for each tract and infant: FA, MD, axial diffusivity (AD), radial diffusivity (RD), QA, normalized QA (nQA), ISO, RDI, normalized RDI at L = 0.6 (nRDI06) and the Jacobian determinant (JDET).

**Fig 2.**
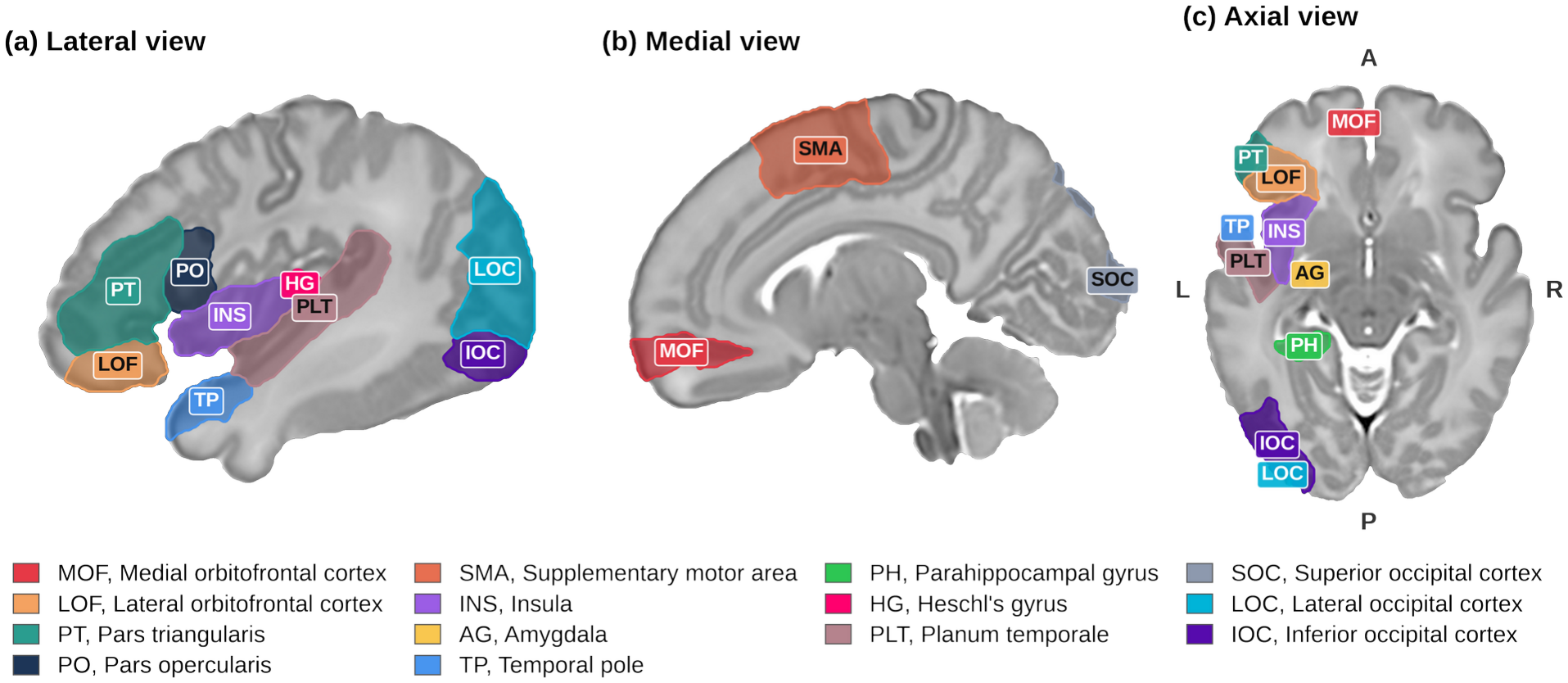
Brain regions associated with language processing. For simplicity, all regions are drawn only on the left hemisphere. Regions are restricted to brain tissue, excluding cerebrospinal fluid; in (b), they are projected over a 4-mm slab lateral to the displayed slice. The cerebrospinal fluid surrounding the brain is masked from the template. Neonate AAL atlas on dHCP group-average T2W template (DSI Studio).

### 2.3 Statistical Analysis

The metrics exported from DSI Studio were analyzed in Python with the statsmodels, pingouin and SciPy libraries. All metric–predictor relationships were modeled with a generalized linear model (GLM) [50]. A single model class was used because a GLM accommodates responses with different distributional support through the choice of family and link function, and it reduces to ordinary least squares for approximately Gaussian responses [50].

The family of each metric was selected by the Akaike information criterion (AIC) [51] on the metric’s distribution pooled across tracts and subjects, and was then fixed for that metric. Near-symmetric diffusivities (MD, AD, RD, ISO, and RDI) were modeled using a Gaussian distribution with an identity link, whereas QA and JDET were modeled using a Gamma distribution with a log link. FA, nQA and nRDI06, which are ratios bounded in the open interval (0, 1) [20], [52], were modeled with beta regression and a logit link [53]. Count-type variables such as streamline count would require a Poisson family; however, they were uninformative and were therefore excluded. Consequently, all retained response variables were continuous. For each metric and tract, the age term was fitted in two forms, linear and quadratic, and the better form was selected by AIC [51], allowing each tract its own developmental trajectory.

Model significance was assessed with a likelihood-ratio test against the null model [54], and the age effect was also tested with the Wald test of its coefficient. Explained variance is reported as the Cox–Snell pseudo-R^2^. The direction of an age effect was taken from the fitted curve, as the sign of the change between the youngest and oldest scan ages. Because many candidate models were fitted across tracts, metrics, predictor sets and functional forms, p-values were corrected with the Benjamini–Hochberg false-discovery-rate (FDR) procedure (α = 0.05) [55] within each tract, hemisphere and predictor group, and only results that survived correction are reported. A model was considered valid when its fit converged. The Durbin– Watson test was not applied, because it assesses autocorrelation among sequentially ordered residuals and is not meaningful for cross-sectional data in which each subject contributes one observation per model [56].

Only infants with complete records for the five demographic and scan variables (scan age, birth age, sex, birth weight and head circumference) were analyzed. Each model contained at most two predictors. All one- and two-predictor combinations, additive and, for pairs, with an interaction term, were fitted with both forms of the age term. This gave 72 candidate models per metric, tract and hemisphere, or 720 per tract and hemisphere for the ten metrics. Each candidate was evaluated by its likelihood-ratio p-value, AIC, Bayesian information criterion (BIC) and the slope of the fitted term. Among the converged fits for a metric, the form of the age term was selected by AIC before significance testing, which kept model selection separate from the test; the p-values were then FDR-corrected.

### 2.4 Age Development Examination

The effect of scan age on each metric was modeled with the GLM described above, with the family fixed per metric. The left and right instances of each tract were modeled separately, and repeated reconstructions and scans had been removed (Section 2.1), so each infant contributed one observation per model. The observations were therefore independent, and a mixed-effects model was not required; with one observation per subject, such a model reduces to the fixed-effects fit. Scan age was entered as a continuous predictor. The other demographic and scan variables (sex, birth age, birth weight and head circumference) were added individually and in pairs to identify the predictors that best explained each metric.

Sex differences were estimated from the additive model of sex and scan age, with female infants as the reference level. Sex was tested by the Wald test of its coefficient (FDR-corrected within each tract and hemisphere), and its effect size was the pseudo-R^2^ gained over the age-only model. The model of sex and head circumference served as a sensitivity analysis.

To resolve development at a finer temporal scale, scan age was also binned into one-week brackets (37–38, 38–39, …, 43–44 weeks; an infant scanned at 37.3 weeks falls into 37–38), and each pair of adjacent brackets was contrasted for every tract and metric. Each contrast was estimated in the family used for that metric in the age model: beta regression for the bounded indices, a Gamma model with log link for the strictly positive metrics, and a Gaussian model for the near-symmetric diffusivities. For each transition we report the likelihood-based p-value, FDR-corrected across the six consecutive-bracket comparisons, and Cohen’s d [57], [58] as a family-independent effect size. This separates when a metric changes within the term window (which transitions are significant) from how much it changes (the effect size).

### 2.5 Reconstruction inconsistency in the 42–43-week bracket

Nearly all infants in the 42–43-week bracket had been reconstructed with a diffusion sampling length ratio of 1.0 instead of 1.25. This shifts the raw QSDR indices (RDI, nRDI06 and QA) downward in that bracket, whereas the tensor metrics, nQA and JDET are not affected. The continuous age model does not absorb this shift. In a linear age model, RDI was significantly associated with PMA in 20 of 22 tracts with all infants but in only 2 of 22 without the 42–43-week bracket; QA rose from 13 to 20 tracts, and nRDI06 was nearly unchanged (22 and 21 tracts). We therefore report the age and sex models of RDI, nRDI06 and QA from the full modeling sweep repeated without the 42–43-week bracket (40 infants), keep all infants for the other seven metrics, and exclude RDI, nRDI06 and QA from the week-resolved analysis. The left–right comparison is within-subject, so a shift common to both hemispheres of an infant does not bias it, and all infants were kept in that analysis.

### 2.6 Laterality

Hemispheric asymmetry was assessed by pairing the homologous left and right tracts of each infant, after outliers had been removed with a robust location–scale criterion. This is the only within-subject contrast in the study, since each infant contributes both values of a tract, and it was therefore analyzed with a paired test rather than with the GLM used for the age and demographic analyses. Normality of the left ™ right differences was assessed with D’Agostino’s test, or with the Shapiro–Wilk test for fewer than 20 pairs (α = 0.05). Normally distributed differences were tested with a paired t-test and the others with the Wilcoxon signed-rank test. A Bonferroni correction was applied across the 27 metrics tested per tract (p < 0.05/27 ≈ 0.0019), and a significant difference was taken to indicate a hemispheric asymmetry in that metric.

## 3. Result

### 3.1 Age Analysis

The expected direction of change, rising anisotropy and falling diffusivity with age, follows the documented course of neonatal white-matter development [5], [19], [59]. MD and RD decreased significantly with age in all 22 tracts (smallest p of order 10^−51^ for RD and 10^−41^ for MD), consistent with ongoing myelination. The strongest associations were in the inferior fronto-occipital fasciculus (RD: pseudo-R^2^ = 0.54 in both IFOF L and IFOF R; MD: 0.49 in IFOF L), so this pathway showed the strongest age dependence of all tracts. FA increased with age in all 22 tracts (pseudo-R^2^ up to 0.42 in IFOF L; smallest p of order 10^−35^), consistent with increasing directional coherence of the axons.

AD changed little: it was significant in only 6 tracts, with pseudo-R^2^ ≤ 0.06, and is therefore a less sensitive marker of neonatal maturation. Without the 42–43-week bracket, RDI decreased in only 8 tracts, whereas nRDI06 decreased in all 22 and QA increased in 21. nQA increased in 20 tracts, most strongly in the FAT, and ISO decreased in 13. JDET increased with age in every tract (pseudo-R^2^ 0.46 in AF L, 0.42 in EC R and 0.53 in UF L and UF R), consistent with volumetric growth of all tracts over the 37–44-week window. Tracts and hemispheres differed in the magnitude of this growth, not in its direction.

A quadratic age term was selected by AIC in 85 of the 178 significant tract–metric cells, including all QA, nRDI06 and RDI cells and 14 FA cells, whereas every MD, RD and JDET cell was best described by a linear trend. Most diffusivity effects remained significant well below the 0.05 threshold after FDR correction. As summarized in Table I, MD, RD and FA were the most informative indicators of microstructural development in this window, with the largest age-related changes in ventral pathways such as the IFOF and ILF.

**TABLE I.** Age-Related Change in Diffusion Metrics of the Language Tracts (37–44 Weeks PMA)

| Metric | AF |  | EC |  | FAT |  | IFOF |  | ILF |  | MLF |  | SLF I |  | SLF II |  | SLF III |  | UF |  | VOF |  |
|---|---|---|---|---|---|---|---|---|---|---|---|---|---|---|---|---|---|---|---|---|---|---|
|  | L | R | L | R | L | R | L | R | L | R | L | R | L | R | L | R | L | R | L | R | L | R |
| **FA** | 0.13°° | 0.06° | 0.27°° | 0.31°° | 0.35°° | 0.37°° | 0.42°° | 0.40° | 0.36° | 0.37° | 0.18°° | 0.30°° | 0.19°° | 0.23°° | 0.26°° | 0.21°° | 0.16°° | 0.17°° | 0.25° | 0.14° | 0.24° | 0.32° |
| **MD** | 0.10° | 0.04° | 0.16° | 0.26° | 0.30° | 0.32° | 0.49° | 0.46° | 0.13° | 0.34° | 0.10° | 0.15° | 0.12° | 0.21° | 0.17° | 0.12° | 0.08° | 0.08° | 0.26° | 0.11° | 0.15° | 0.17° |
| **RD** | 0.12° | 0.04° | 0.19° | 0.30° | 0.37° | 0.40° | 0.54° | 0.54° | 0.27° | 0.41° | 0.13° | 0.23° | 0.16° | 0.26° | 0.23° | 0.17° | 0.11° | 0.12° | 0.33° | 0.15° | 0.21° | 0.25° |
| **JDET** | 0.46° | 0.30° | 0.32° | 0.42° | 0.06° | 0.18° | 0.32° | 0.38° | 0.04° | 0.25° | 0.27° | 0.35° | 0.23° | 0.36° | 0.08° | 0.15° | 0.21° | 0.35° | 0.53° | 0.53° | 0.19° | 0.16° |
| **QA** | 0.03°° | 0.06°° | 0.11°° | 0.11°° | 0.18°° | 0.23°° | 0.10°° | 0.12°° | 0.21°° | 0.12°° | 0.07°° | 0.13°° | 0.07°° | 0.11°° | 0.08°° | 0.09°° | 0.07°° | 0.05°° | 0.06°° | 0.01° | 0.06°° | 0.10°° |
| **nQA** | 0.00° | 0.01° | 0.05°° | 0.13° | 0.24° | 0.28° | 0.16° | 0.20°° | 0.21° | 0.21°° | 0.06° | 0.12°° | 0.05° | 0.10° | 0.07° | 0.08° | 0.02°° | 0.02° | 0.09°° | 0.04°° | 0.09°° | 0.16°° |
| **AD** | 0.01° | 0.04°° | 0.01°° | 0.03° | 0.01° | 0.00° | 0.02° | 0.03°° | 0.06° | 0.00° | 0.00° | 0.01° | 0.00° | 0.00° | 0.00° | 0.01°° | 0.00° | 0.02°° | 0.06° | 0.02° | 0.00° | 0.01°° |
| **RDI** | 0.03°° | 0.04°° | 0.06°° | 0.02°° | 0.01°° | 0.02°° | 0.02°° | 0.00° | 0.00° | 0.00° | 0.02°° | 0.06°° | 0.03°° | 0.02°° | 0.02°° | 0.03°° | 0.05°° | 0.05°° | 0.01°° | 0.01° | 0.01° | 0.00° |
| **ISO** | 0.04°° | 0.03°° | 0.06°° | 0.03°° | 0.02°° | 0.01°° | 0.02°° | 0.02°° | 0.02° | 0.02°° | 0.04°° | 0.05°° | 0.03°° | 0.02°° | 0.04°° | 0.03°° | 0.06°° | 0.06°° | 0.02°° | 0.02° | 0.03° | 0.00° |
| **nRDI06** | 0.11°° | 0.04°° | 0.14°° | 0.18°° | 0.12°° | 0.12°° | 0.14°° | 0.15°° | 0.11°° | 0.14°° | 0.12°° | 0.11°° | 0.09°° | 0.11°° | 0.12°° | 0.09°° | 0.12°° | 0.11°° | 0.12°° | 0.10°° | 0.13°° | 0.09°° |

increase decrease darker = smaller p (p < 10<sup>−6</sup>, < 10<sup>−3</sup>, < 0.01, < 0.05) ° linear °° quadratic
Rows are metrics and columns are tracts by hemisphere (L, left; R, right). Each cell is one GLM of the metric on PMA at scan, with the family matched to the metric's support (Beta/logit: FA, nQA, nRDI06; Gamma/log: QA, JDET; Gaussian/identity: MD, RD, AD, ISO, RDI); the number is the Cox–Snell pseudo-R <sup>2</sup> and the symbol the AIC-selected form of the age term (° linear, °° quadratic). Red shading marks an increase and blue a decrease between the youngest and oldest scan ages; darker shading marks a smaller Benjamini–Hochberg-corrected p-value (bands p < 10 <sup>−6</sup>, < 10<sup>−3</sup>, < 0.01, < 0.05). Values in grey without shading were not significant after correction. QA, RDI and nRDI06 are estimated without the 42–43-week bracket (see the section on the 42–43-week bracket).

### 3.2 Weekly Age Analysis

Each pair of consecutive one-week brackets (37–38 → 38–39, …, 42–43 → 43–44) was contrasted for every metric within the metric’s GLM family (Section 2.4), with FDR correction across the six transitions. Six metrics (FA, JDET, MD, nQA, RD and AD) showed significant week-to-week change in at least one tract (Supplementary Table S3), although AD did so in only two transitions. ISO showed no significant weekly transition, consistent with a gradual change that is not concentrated in any single week. QA, RDI and nRDI06 were not tested because of the reconstruction inconsistency in the 42–43-week bracket (Section 2.5).

The week-resolved analysis (Table II) showed that anisotropy and volume changed at different times. All 35 significant FA transitions were increases; they peaked at 38–39 → 39–40 weeks (12 tracts), and none occurred at the final transition. JDET transitions continued to the end of the window: all 45 were increases, 20 of them at 42–43 → 43–44, with the largest effects in the left IFOF (d = +1.53) and the left UF (d = +1.48) at that transition. All diffusivity transitions were decreases. MD transitions (24) were spread across the window, whereas RD transitions (31) were absent at the final transition. This pattern of early anisotropy change and volumetric growth continuing to the end of the term window is averaged out by the continuous age model (Table I) and becomes visible only at weekly resolution.

**TABLE II.**
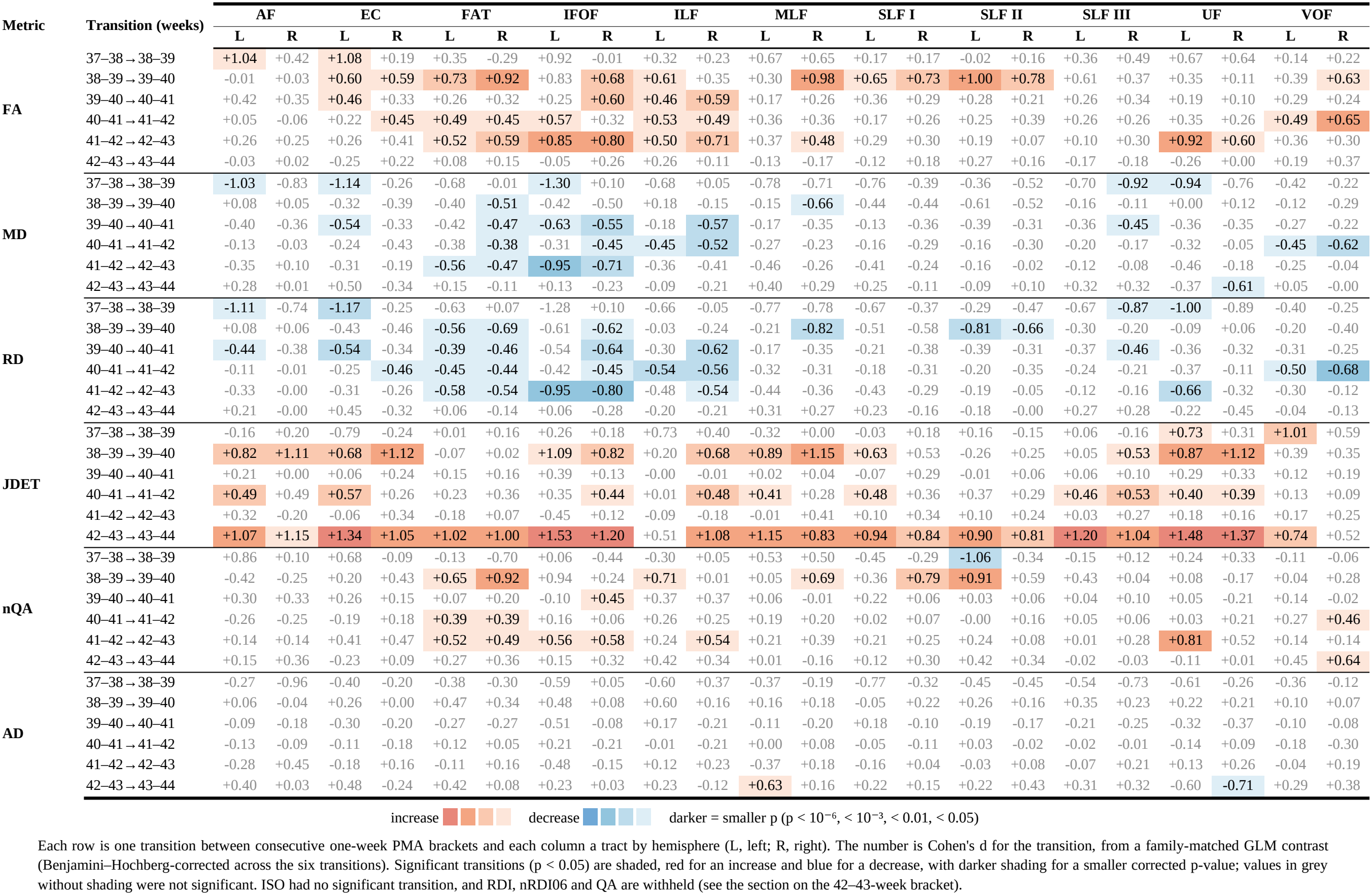
Week-Resolved Development of the Language Tracts (37–44 Weeks PMA)

### 3.3 Laterality Analysis

Paired tests with Bonferroni correction showed significant left–right differences in 86 of the 110 tract– metric pairs, 54 favoring the left hemisphere and 32 the right (Table III). Leftward asymmetry predominated in the classical language pathways: the AF (6 leftward vs 1 rightward metric), the IFOF (6 vs 1), the MLF (7 vs 0) and the second branch of the superior longitudinal fasciculus (SLF II: 7 vs 2). In these tracts, the left hemisphere had higher FA and lower MD and RD. Rightward asymmetry characterized the UF (0 vs 8) and was also present in the ILF (4 vs 5) and parts of the FAT. The core language pathways are therefore already lateralized toward the left hemisphere in term neonates, which is consistent with an early onset of left-hemisphere specialization for language.

**TABLE II.** Hemispheric Asymmetry of Diffusion Metrics in the Language Tracts (37–44 Weeks PMA)

| **Metric** | **AF** | **EC** | **FAT** | **IFOF** | **ILF** | **MLF** | **SLF I** | **SLF II** | **SLF III** | **UF** | **VOF** |
|---|---|---|---|---|---|---|---|---|---|---|---|
| **FA** | L | L | L | L | R | L | R |  |  | R | R |
| **MD** | L | L | L |  | R | L | L | L |  | R |  |
| **RD** | L | L | L |  | R | L | L | L |  | R | R |
| **JDET** | R | R | R |  | L | L |  | L | L |  | L |
| **QA** |  | L | L | L | R |  | R | R |  | R | R |
| **nQA** |  | L | L | L | R |  | R | R |  | R | R |
| **AD** |  | L | R | R |  | L | L | L |  |  | L |
| **RDI** | L | R | R | L | L |  | R | L | L | R | L |
| **ISO** | L |  | R | L | L | L | R | L | L | R | L |
| **nRDI06** | L | L |  | L | L | L |  | L | L | R | L |
| **Leftward (n)** | **6** | **7** | **5** | **6** | **4** | **7** | **3** | **7** | **4** | **0** | **5** |
| **Rightward (n)** | **1** | **2** | **4** | **1** | **5** | **0** | **5** | **2** | **0** | **8** | **4** |

L leftward R rightward blank = no asymmetry
L marks a significant asymmetry favoring the left hemisphere and R one favoring the right, where the favored hemisphere is the one with the more mature value (higher FA, QA, nQA and JDET; lower MD, RD, AD, ISO, RDI and nRDI06); blank cells showed no asymmetry (paired t-test, or Wilcoxon signed-rank test where normality of the differences was rejected; Bonferroni-corrected, p < 0.05/27 ≈ 0.0019). The bottom rows count leftward and rightward asymmetries per tract. Morphological measures are analysed in the companion article and are not counted here.

### 3.4 Gender Analysis

After adjustment for scan age, sex differences were found in all 22 tracts, and all were small (Table IV). JDET was the strongest and most consistent sex-related metric. It was larger in males in every tract (pseudo-R^2^ gain 0.03–0.14), most strongly in FAT L (0.14, p = 3.3 × 10^−9^). MD (3 tracts) and RD (4 tracts) were higher in males, FA was higher in females in three tracts (FAT R, ILF L and VOF R), and QA and nQA showed no sex difference. The left ILF showed both a JDET effect favoring males (0.13) and an FA effect favoring females (0.02). When head circumference instead of age was adjusted for, the JDET difference remained in only 7 tracts, whereas higher FA (10 tracts) and nQA (5) in females and higher MD (9) and RD (10) in males became more widespread. Overall, sex had only a small and non-uniform effect on the microstructure of these tracts.

**TABLE IV.** Sex Differences in Diffusion Metrics of the Language Tracts, Adjusted for PMA at Scan.

| Metric | AF |  | EC |  | FAT |  | IFOF |  | ILF |  | MLF |  | SLF I |  | SLF II |  | SLF III |  | UF |  | VOF |  |
|---|---|---|---|---|---|---|---|---|---|---|---|---|---|---|---|---|---|---|---|---|---|---|
|  | L | R | L | R | L | R | L | R | L | R | L | R | L | R | L | R | L | R | L | R | L | R |
| **FA** | .00 | .02 | .00 | .00 | .01 | .02 | .00 | .00 | .02 | .01 | .00 | .00 | .02 | .01 | .01 | .01 | .00 | .00 | .01 | .00 | .01 | .03 |
| **MD** | .01 | .03 | .06 | .00 | .02 | .02 | .01 | .00 | .01 | .01 | .01 | .04 | .03 | .03 | .02 | .04 | .04 | .04 | .01 | .00 | .01 | .04 |
| **RD** | .01 | .03 | .07 | .00 | .02 | .02 | .01 | .00 | .02 | .01 | .01 | .05 | .04 | .03 | .03 | .04 | .04 | .04 | .01 | .00 | .02 | .04 |
| **JDET** | .07 | .13 | .07 | .03 | .14 | .11 | .06 | .04 | .13 | .08 | .06 | .05 | .05 | .05 | .10 | .10 | .08 | .05 | .07 | .05 | .09 | .12 |
| **QA** | .02 | .04 | .06 | .02 | .02 | .03 | .02 | .01 | .02 | .00 | .03 | .06 | .02 | .02 | .03 | .03 | .04 | .02 | .01 | .00 | .01 | .01 |
| **nQA** | .01 | .02 | .01 | .00 | .01 | .01 | .00 | .01 | .01 | .00 | .01 | .00 | .00 | .00 | .01 | .00 | .00 | .00 | .00 | .00 | .01 | .01 |
| **AD** | .00 | .01 | .00 | .00 | .00 | .00 | .01 | .00 | .00 | .00 | .00 | .01 | .00 | .01 | .00 | .00 | .00 | .00 | .00 | .00 | .00 | .01 |
| **RDI** | .01 | .06 | .02 | .01 | .00 | .00 | .00 | .00 | .00 | .00 | .01 | .04 | .01 | .01 | .01 | .01 | .01 | .02 | .00 | .00 | .00 | .00 |
| **ISO** | .00 | .00 | .00 | .00 | .00 | .00 | .00 | .00 | .00 | .00 | .01 | .00 | .00 | .00 | .00 | .00 | .00 | .00 | .00 | .00 | .00 | .00 |
| **nRDI06** | .00 | .05 | .00 | .02 | .01 | .01 | .01 | .02 | .02 | .02 | .01 | .02 | .00 | .01 | .01 | .00 | .01 | .01 | .01 | .01 | .02 | .01 |

higher in males higher in females darker = smaller p (p < 10<sup>−6</sup>, < 10<sup>−3</sup>, < 0.01, < 0.05)
Each cell is a GLM of the metric on sex and PMA (family as in Table I; female infants are the reference level), with sex tested by the Wald test of its coefficient. The number is the Cox–Snell pseudo-R<sup>2</sup> gained over the age model; red shading marks a higher value in males and blue in females at the mean PMA, and darker shading a smaller Benjamini–Hochberg-corrected p-value. Values in grey without shading were not significant after correction.

## 4. Discussion

This study examined the microstructural development of the language pathways in term neonates, focusing on how diffusion metrics change over the first weeks after birth. Using a large sample (n = 310) and both tensor and QSDR reconstructions, we derived tract-specific metrics for the major perisylvian fibers implicated in early language function. Below we discuss the implications of these findings for neonatal neurodevelopment, the limitations of the study, and directions for future research.

All examined language tracts showed significant changes in diffusion metrics across the neonatal period (37–44 weeks PMA), underlining the importance of this short window for white-matter maturation. MD and RD decreased, and FA increased with age, indicating increasing microstructural organization. These observations agree with previous reports of rapid fiber development during infancy, particularly in the first months of life [59], [60]. In densely sampled infants, white-matter growth at birth is several times faster than at six months [61]. JDET increased in every tract, indicating that the language pathways grow in volume throughout the term period.

Some pathways, including the FAT and the right IFOF and ILF, showed more week-to-week changes than others, which shows the value of fine temporal resolution in neonatal neuroimaging. These tract-specific trajectories support the concept of asynchronous maturation within the language network [7], [62]. In dHCP neonates, the coupling between white-matter structure and function also develops heterogeneously across the major fiber tracts, with a left-lateralized tendency in the ILF and IFOF [63].

Our data are consistent with the current multi-tract view of the language network [12], [64]: several white-matter tracts, not the arcuate fasciculus alone, showed marked microstructural growth in the neonatal period. Ventral pathways, such as the IFOF and UF, often mature earlier in infancy than dorsal pathways [7], whereas dorsal pathways, such as the AF and SLF, continue to mature over a longer period; the neurite density of dorsal language tracts is relatively low near birth and rises steeply afterwards [65]. Consistent with these observations, the rate of microstructural change differed between tracts in our cohort. The tract-specific weekly patterns, especially in the FAT and IFOF, are in line with reports of heterogeneous maturation between and within white-matter pathways [66], [67].

Leftward asymmetries were found in classical language tracts such as the AF and IFOF, consistent with earlier reports of a leftward asymmetry of the infant AF [68] and with an emerging left-hemisphere specialization for language. Structural asymmetry of the language network emerges early: perisylvian regions are already asymmetric in preterm newborns [27], and the macrostructural asymmetry of the arcuate fasciculus is present by two years of age [69]. The leftward asymmetry we found in FA and diffusivity, for example in the AF, is consistent with the structural asymmetries of perisylvian regions and of the dorsal pathway reported in newborns and young infants [27]–[29], [62]. A leftward IFOF fiber cross-section has also been reported in preterm-born infants followed to six months, although the same study found a rightward FA asymmetry in language-processing regions [70]. This structural asymmetry may be a developmental substrate of the left-hemisphere dominance for language, which is present before substantial postnatal language exposure [71].

However, the evidence on early white-matter lateralization is not uniform. In infants aged 0–24 months, the AF and UF matured at similar rates in both hemispheres, and their left lateralization was not present in infancy [72]. In the first postnatal months, the lateralization of the major tracts also varies with age and between tracts [31]. These findings do not necessarily conflict with ours, because we compared the level of each metric between hemispheres within a narrow window around term, whereas [72] compared rates of change over two years.

Among the variables examined (GA at birth, birth weight and head circumference), age at scan was the strongest predictor of the microstructural metrics across language tracts. Earlier newborn studies likewise found gestational age to be the birth indicator most strongly associated with white-matter maturation [38], and GA at birth to be more strongly associated with FA than postnatal age [32]. PMA at scan combines both, so within the 37–44-week window, myelination and fiber organization appear to follow maturational age most closely. Sex differences were small, and the larger tract volume in males was largely accounted for by head size, which is larger in male newborns [73]. This agrees with reviews indicating that sex differences in brain structure or function do not necessarily translate into differences in language outcome [36]. Taken together, these results show that the dorsal and ventral language tracts of term neonates undergo rapid growth and microstructural refinement, which may provide a structural basis for later linguistic and cognitive abilities.

Several methodological factors limit our conclusions. Although the sample was large (n = 310) and covered 37–44 weeks, the data are cross-sectional, so individual growth trajectories cannot be confirmed. Longitudinal studies [74] show that some microstructural changes follow nonlinear trajectories, which cross-sectional data can only approximate. In addition, the cohort is not uniformly distributed over 37–44 weeks: the youngest and oldest brackets contain fewer infants (Supplementary Fig. S1), which reduces the precision of the week-resolved contrasts at the ends of the window. The large number of tests across tracts, metrics, and predictors required correction for multiple comparisons. Although the false-discovery rate was controlled, a larger sample would reduce both type I and type II errors. Finally, the 42–43-week bracket was reconstructed with a different sampling length ratio, which affects RDI, nRDI06 and QA. These metrics were therefore estimated without that bracket, and the affected infants should be reconstructed again with uniform parameters before these indices are reported for the full cohort.

QSDR reconstruction mitigates some limitations of tensor-based tractography, such as crossing fibers [25], and improves the accuracy of fiber reconstruction [47]. Partial-volume effects remain a concern, however, especially in the largely unmyelinated neonatal brain. Some pathways were difficult to reconstruct in every infant, notably the right AF (156 of 310 infants) and the left IFOF (207). The low reconstruction rate of the right AF agrees with the strong leftward volumetric asymmetry of the AF in adults [75]. In adults, AF laterality estimates depend strongly on the tractography algorithm [76], and reconstructions that resolve crossing fibers remain uncommon in infant tractography [77], which limits direct comparison with earlier tensor-based studies. Tractography also relied on an adult-derived atlas (HCP842). Differences in neonatal brain morphology can cause misalignment even after nonlinear registration; atlases that place neonates and adults in a common space have since become available [78]. Although pruning and visual verification were applied, small local misregistrations could affect the microstructural measures. Future pipelines could use stronger diffusion gradients, more advanced diffusion models (e.g., NODDI or DKI, which has been applied in neonates [79]) and improved fiber dissection to separate adjacent tracts with overlapping cortical endpoints. This is particularly relevant for neonatal pathways in border zones, such as the extreme capsule and the middle longitudinal fasciculus.

Although we discuss how structural asymmetry may underlie language lateralization, the study did not include functional imaging or behavioral measures of language. Neonatal white-matter maturation is associated with language development at 12 months [34], and myelination at 7 months predicts later language [80], but testing this relationship in the same cohort requires longitudinal neurodevelopmental assessment. Repeated imaging combined with behavioral or electrophysiological assessment of language, such as auditory event-related potentials, would clarify how structural changes relate to developing language skills, but such multimodal data remain sparse. In infancy and childhood, more adult–child conversation and greater language input are associated with stronger white-matter connectivity and myelination [81], [82]. In preterm neonates, increased exposure to maternal speech was followed by more mature microstructure of the left AF [83], whereas greater language input in infancy has also been linked to lower anisotropy of the left posterior AF at two years [84]. Whether such inputs modulate neonatal microstructure, particularly in the dorsal or ventral pathways, remains to be tested.

Neonates with known pathologies, severe motion artifacts or incomplete anthropometric data were excluded from the study. Future studies should therefore investigate whether the present findings extend to more diverse neonatal populations, particularly preterm neonates (< 37 weeks), neonates with perinatal complications and those with atypical growth and development.

By charting tract-specific development and hemispheric asymmetries, this work provides a detailed map of the perisylvian language pathways in healthy term neonates. Such normative data could serve as a reference for detecting deviations associated with language or cognitive impairment in high-risk cohorts, such as very preterm infants or infants at genetic risk of neurodevelopmental disorders. Combining neonatal imaging with predictive modeling may improve the early identification of infants at risk for dyslexia, developmental language disorder, or autism spectrum disorder. White-matter connectomes at birth have predicted cognitive scores at two years [85], although an attempt to predict later outcomes from neonatal cortical microstructure in dHCP infants did not exceed chance [86]. Knowing which tracts deviate earliest could inform screening tools or interventions that support language acquisition.

## 5. Conclusion

By integrating advanced diffusion imaging techniques, an unprecedented sample size, and focused analyses of perisylvian language pathways, our work underscores the remarkable neuroplasticity and organizational specificity inherent in the newborn brain. This study thus provides an essential step toward understanding how the earliest foundations of language networks set the stage for lifelong communication and cognitive development. We demonstrated that language-related white matter tracts exhibit discernible and tract-specific microstructural growth in the first weeks of neonatal life. These changes are strongly associated with chronological age and show distinct weekly differences in certain pathways, and often align with left-lateralized organization. Our results reinforce that structural asymmetry emerges early, potentially scaffolding the rapid acquisition of language functions in infancy. Future longitudinal, multi-modal studies will be crucial for linking these neonatal structural trajectories to subsequent cognitive and linguistic outcomes, ultimately contributing to refined clinical benchmarks and interventions for at-risk pediatric populations.

## Declarations

## Data and code availability

The dHCP data are publicly available from the developing Human Connectome Project (www.developingconnectome.org) under its data-use terms. The analysis code is available at https://github.com/alikaz3mi/tract_analysis. Derived tract metrics are available from the corresponding author on request.

## Author contributions

A. Kazemi: conceptualization, methodology, software, formal analysis, investigation, visualization, writing – original draft. R. Namiranian: methodology, validation, writing – review and editing. H. Abrishami Moghaddam: conceptualization, methodology, validation, supervision, project administration, writing – review and editing.

## Ethics

This study used only publicly released, de-identified dHCP data; the dHCP obtained ethical approval and parental consent.

## Acknowledgments

Data were provided by the developing Human Connectome Project, KCL-Imperial-Oxford Consortium, funded by the European Research Council under the European Union Seventh Framework Programme (FP/2007-2013) / ERC Grant Agreement no. 319456. We are grateful to the families who generously supported this trial.

